# CD24-Fc resolves inflammation and rescues CD8 T cells with polyfunctionality in humanized mice infected with HIV-1 under cART

**DOI:** 10.1101/2024.12.16.628615

**Authors:** Guangming Li, Jianping Ma, Haisheng Yu, Ourania Tsahouridis, Yaoxian Lou, Xiuting He, Masaya Funaki, Poonam Mathur, Shyamasundaran Kottilil, Pan Zheng, Yang Liu, Lishan Su

## Abstract

The persistence of HIV-1 reservoirs during combination anti-retroviral therapy (cART) leads to chronic immune activation and systemic inflammation in people with HIV (PWH), associating with a suboptimal immune reconstitution as well as an increased risk of non-AIDS events. This highlights the needs to develop novel therapy for HIV-1 related diseases in PWH. In this study, we assessed the therapeutic effect of CD24-Fc, a fusion protein with anti-inflammatory properties that interacts with danger-associated molecular patterns (DAMPs) and siglec-10, in chronic HIV-1 infection model using humanized mice undergoing suppressive cART. Our findings show that CD24-Fc treatment significantly reduced inflammation and immune hyperactivation in vivo when combined with cART. CD24-Fc mediated resolution of inflammation was associated with improved recovery of CD4 T cells, reduced immune activation, restored central memory T cells and reversal of immune cell exhaustion phenotype. Notably, CD24-Fc treatment rescued CXCR5+ CD8 central memory T cell (T_CM_) which correlated with increased polyfunctionality in HIV-specific T cells in humanized mice and in cultured peripheral blood mononuclear cells (PBMCs) from PWH. This restoration of CXCR5+ memory CD8 T cells was associated with HIV replication inhibition, delayed viral rebound and reduced HIV-1 pathogenesis upon cART cessation. This study suggests that CD24-Fc treatment could represent a promising new therapeutic strategy for managing chronic systemic inflammation and associated diseases in PWH.

**Author summary:** Combination antiretroviral therapy (cART) cannot block viral gene expression from activated HIV proviral DNA in reservoir cells, contributing to chronic immune activation and inflammation associated diseases in people with HIV (PWH). The therapeutic treatment of anti-inflammatory fusion protein CD24-Fc in humanized mice during suppressive cART (*i*) resolves inflammation and chronic HIV-1 immune pathogenesis during suppressive cART, (*ii*) rescues CXCR5-expressing CD8 memory T cells and enhances antiviral response in humanized mice and PWH PBMCs, (*iii*) delays virus rebound and reduces viral pathogenesis after cART cessation. Thus, CD24-Fc could provide a novel therapeutic strategy for treating chronic systemic inflammation and associated diseases in PWH.

## Introduction

The primary barrier to an HIV cure is the persistence of the HIV-1 reservoir during combination antiretroviral therapy (cART). Modern cART is a highly effective treatment that enables people with HIV (PWH) to achieve undetectable viral loads, preventing HIV transmission [1, 2]. However, cART requires lifelong adherence due to the presence of cART-resistant reservoirs, which cause rapid viral rebound if treatment is interrupted. HIV-1 reservoir persistence is often accompanied by residual inflammation that supports reservoir stability, and this relationship may be bidirectional [3]. Approximately 20% of individuals who initiate cART with low CD4 counts experience poor CD4 T cell reconstitution, which correlates with a higher risk of non-AIDS-related complications [4–8]. Furthermore, individuals with suboptimal CD4 recovery tend to exhibit greater immune activation and inflammation compared to those with better CD4 recovery [9], while PLWH who continue to experience inflammation despite effective cART are at an elevated risk for comorbidities and non-AIDS events [10–12]. Therefore, residual inflammation and immune activation are believed to play critical roles in HIV-1 associated diseases in post cART era. Targeting residual inflammation in HIV-1 infection may offer a promising therapeutic avenue for managing HIV-1 and related diseases.

Recent studies have explored therapeutic modulation of the inflammatory response in PWH, including anti-inflammatory drugs targeting specific receptors or cytokines, as well as immunomodulatory supplements [13]. Although some treatments reduce immune activation and inflammation associated with HIV-1, none have significantly improved anti-HIV immune responses or reservoir elimination. Using a humanized mouse model, we and others have demonstrated that blocking type I interferon (IFN-I) signaling during cART reduces systemic inflammation and immune activation, enhancing anti-HIV immunity and promoting HIV-1 reservoir clearance [14, 15]. We further showed that blocking IFN-I receptors or depleting IFN-I-producing cells in cART-naïve animals restored human immune cell viability, cell counts, and function [16, 17]. These findings underscore the potential of anti-inflammatory therapies as novel immunotherapies for HIV-1 associated diseases.

HIV-1 infection induces cell death both directly and indirectly through various pathways [18–22], resulting in persistent cell death, immune activation, inflammation, and tissue damage even with cART [9, 23–26]. Inflammatory responses triggered by cell death and tissue damage are well-documented in chronic diseases [27, 28], where danger-associated molecular patterns (DAMPs) released during cellular stress promote inflammation and immune activation [29, 30], play a crucial role in promoting inflammatory response and immune activation during cells death or tissue damage [31]. Blocking DAMP signaling may thus offer a new anti-inflammatory approach for managing chronic HIV-1 disease.

In non-human primates (NHPs), we investigated the therapeutic potential of the human CD24-Fc fusion protein, which mitigates inflammation through interactions with DAMPs and siglec-10. CD24-Fc conferred protection against weight loss, wasting syndrome, intractable diarrhea, and decreased AIDS morbidity and mortality in pathogenic SIV infection [32]. Remarkably, CD24-Fc also lowered the incidence of pneumonia and protected against acute respiratory distress syndrome (ARDS) in these NHPs [32, 33]. Additionally, a recent phase III trial (NCT04317040) demonstrated that CD24-Fc effectively reduced systemic inflammation and promoted immune homeostasis in severe COVID-19 patients, without compromising the anti-viral antibody response [34]. However, the effects of CD24-Fc on HIV-1 reservoir persistence and immune pathogenesis during cART remain unclear. Humanized mice engrafted with human immune cells are valuable for studying HIV-1 infection, pathogenesis, and therapies [15–17, 35–43]. In this study, we evaluate the therapeutic potential of CD24-Fc in a chronic HIV-1 infection model using suppressive cART in humanized mice.

## Results

### CD24-Fc treatment resolves the residual inflammation in chronic HIV-1 infection during suppressive cART

In this preclinical study evaluating CD24-Fc therapy in HIV-1 disease, we used a humanized mouse model infected with the HIV-1 JRCSF strain. Mice began combined antiretroviral therapy (cART) at 4 weeks post-infection (wpi) after chronic infection had been established, and HIV-1 viremia was subsequently suppressed to undetectable levels following 5 weeks of treatment. CD24-Fc therapy was then introduced via intraperitoneal (i.p.) injection twice weekly, starting at 11 wpi, when viremia was stably suppressed for 2 weeks, and continued until 3 days before euthanasia at 15 wpi. Notably, no viremia blip was observed in any infected animals following suppression by cART, indicating effective viral inhibition and no stimulatory effect of CD24-Fc on HIV-1 replication (Fig. 1a). At termination, we measured inflammatory cytokines in blood, including IP-10, IL-10, GM-CSF, MCP-1, and MIP-1b. CD24-Fc treatment completely resolved the residual inflammatory response compared to both the cART + Ig and uninfected groups (Fig. 1b). This reduction in inflammation was further confirmed by significantly decreased levels of interferon-stimulated genes (ISGs) in splenocytes in the CD24-Fc group compared to the cART + Ig group. Additionally, ISG levels in CD24-Fc treated animals were comparable to those in HIV-naïve controls, indicating that CD24-Fc effectively resolved HIV-associated inflammation during suppressive cART (Fig. 1c).

**Figure 1.**
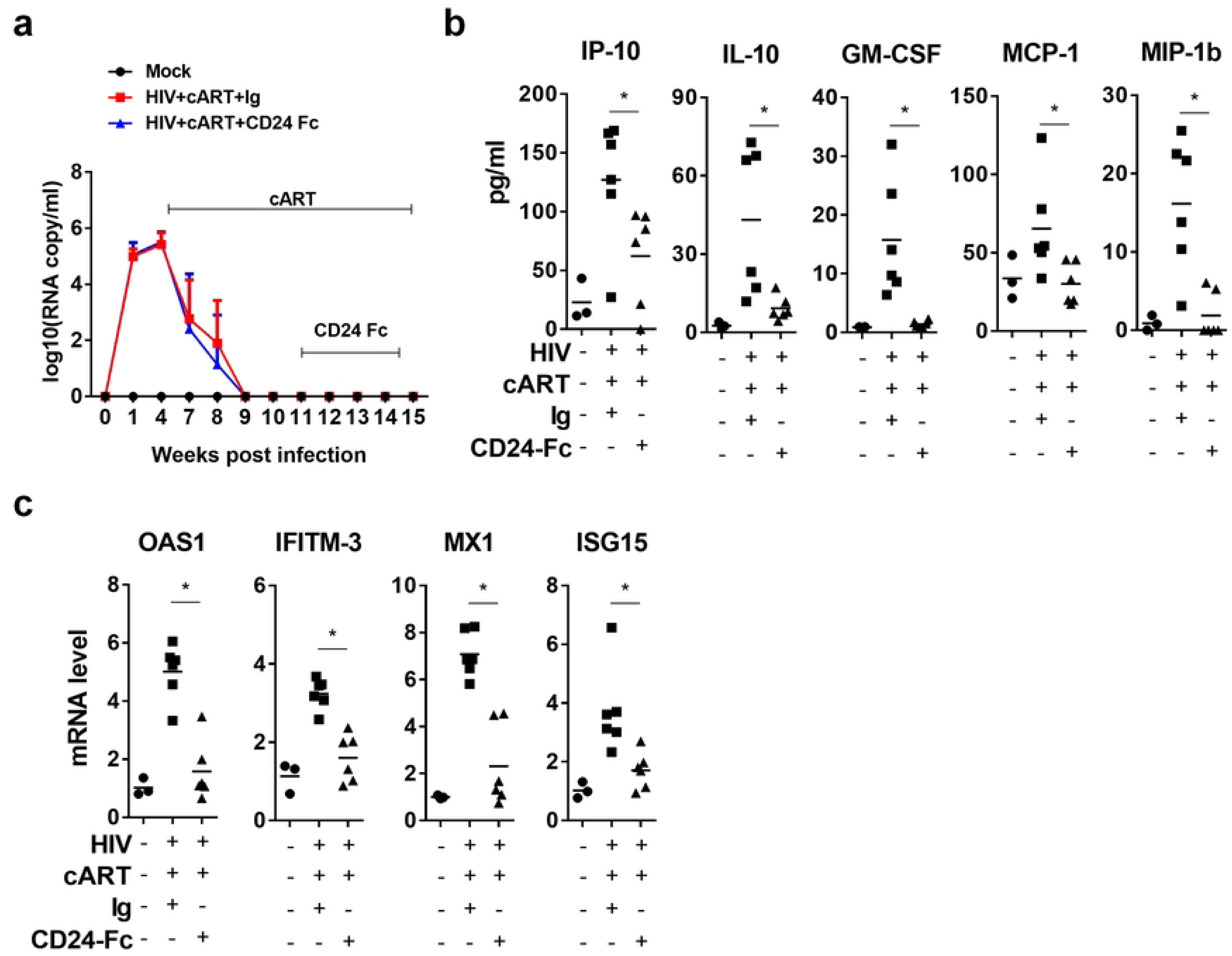
CD24-Fc treatment resolves residual inflammation and reverse HIV immune pathogenesis during cART. Humanized mice were infected and treated as in Fig.1a and terminated at 15 wpi. (a) HIV viral load. (b) Cytokines in blood measured by Luminex. (c) mRNA level of ISGs in splenocytes detect by real-time PCR. Bar represents mean value. * = p<0.05. Error bar in Fig.1a indicates mean value ±s.e.m.

HIV-1 primarily target CD4 T cells, infected and deplete them [44]. cART is able to rescue CD4+ T cells in HIV patients, somehow, the reconstitution of CD4+ T cells fails in about 30% patients [45, 46]. In this study, the cART + Ig group still showed a significantly lower CD4+/CD8+ T cell ratio in the spleen compared to the uninfected (mock) group. In contrast, CD24-Fc treatment restored the CD4+/CD8+ T cell ratio to levels comparable to the mock group (Fig. 2a, Fig. S1a). Although cART effectively suppressed viral replication, infected animals displayed persistent immune activation, evidenced by an elevated frequency of HLA-DR and CD38 double-positive CD8+ T cells in the cART + Ig group (Fig. 2b, Fig. S2b). CD24-Fc treatment, however, significantly reduced immune activation in comparison with both the cART + Ig and mock groups, indicating effective resolution of immune activation (Fig. 2b, Fig. S2b). Chronic HIV-1 infection often results in the depletion of central memory CD8+ T (T_CM_) cells, which do not fully recover despite effective viral suppression [47]. Similarly, we observed that HIV-1 infection in humanized mice led to reduced T_CM_ cell frequency and a lower T_CM_/effector memory T cell (T_EM_) ratio, even with cART-mediated viral suppression (Fig. 2c, Fig. S2c). Remarkably, CD24-Fc treatment in combination with cART restored T_CM_ cell levels and reversed the T_CM_/T_EM_ ratio to that of uninfected animals (Fig. 2c, Fig. S2c). Moreover, HIV infection reduces the proportion of CD28-expressing CD8+ T cells, and this loss is not fully reversible with cART alone. [48–50]. Consistent with human studies, our cART alone showed decreased CD28+ CD8+ T cell frequency and reduced CD28 expression intensity among CD8+ T cells. CD24-Fc treatment with cART, however, restored both CD28+ CD8+ T cell frequency and CD28 expression levels (Fig. 2d-f). These findings suggest that chronic inflammation persists under suppressive cART and that cART alone is insufficient to resolve HIV-1-induced immune pathogenesis. CD24-Fc demonstrated an anti-inflammatory effect, resolving chronic HIV-1-associated immune pathology during cART.

**Figure 2.**
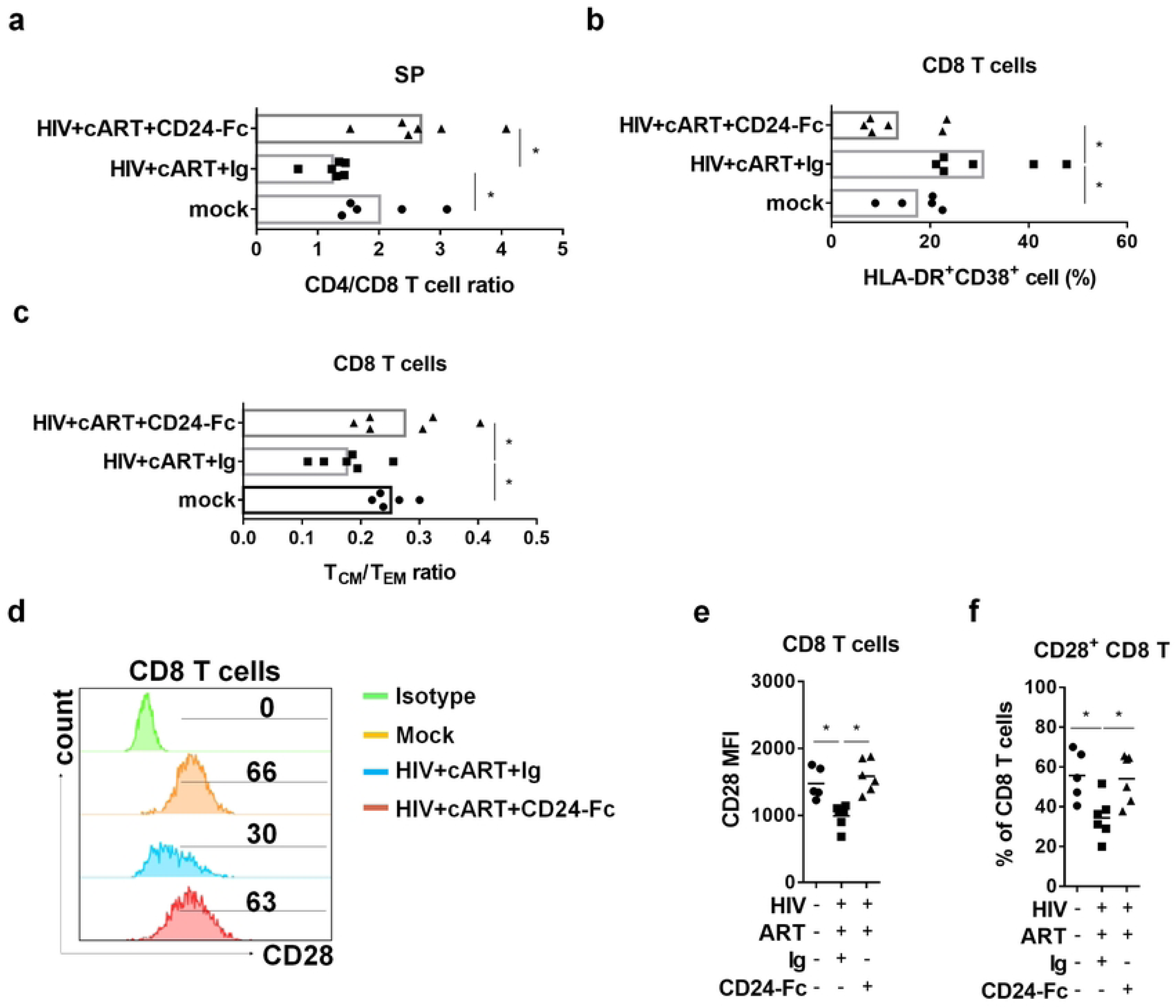
CD24-Fc treatment reverses HIV immune pathogenesis during cART. Humanized mice were infected and treated as in Fig.1a. Splenocytes were analyzed on termination. (a) The ratio of CD4+ T/CD8+ T cell. (b) Summary graph show the frequency of HLA-DR and CD38 double positive CD8+ T cells. (c) The ratio of central memory/effector memory CD8+ T cell. (d) Representative histograms show CD28 expression in CD8 T cells in different group. Bar gates indicate the percentage of CD28+ CD8 T cells. (e) Summary graph for CD28 mean fluorescence intensity (MFI) in CD8 T cells. (f) Summary graph for the percentage of CD28+ CD8 T cells. Bar represents mean value. * = p<0.05.

### CD24-Fc treatment rescues CXCR5+ CD8 T cells and anti-HIV T cell response in vivo

A specific subset of CD8+ T cells expressing the chemokine receptor CXCR5 is crucial for controlling viral replication, with its levels inversely correlated with HIV-1 viral load in PWH [51]. In our study, CD8+ T cells from the cART + Ig group clustered differently in UMAP space compared to both the mock and cART + CD24-Fc groups (Fig. 3a). Specifically, HIV-1-infected animals under suppressive cART showed a significantly reduced frequency of CXCR5+ CD8 T_CM_ cells in the spleen compared to uninfected animals. CD24-Fc treatment reversed this reduction, restoring CXCR5+ T_CM_ levels to those observed in the mock group (Fig. 3b and 3c).

**Figure 3.**
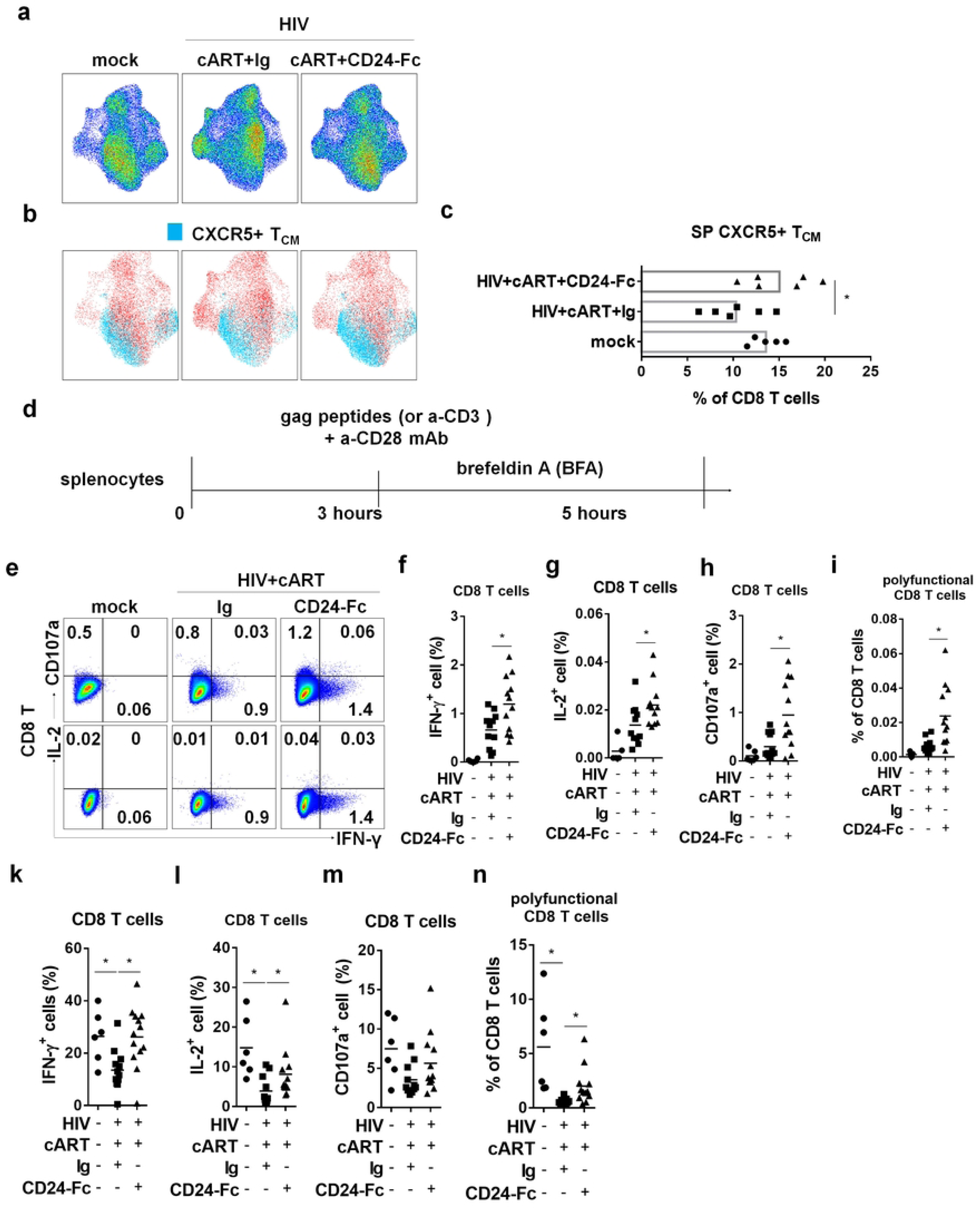
CD24-Fc treatment rescues CXCR5+ CD8 T cells and anti-HIV T cell response in vivo. Humanized mice were infected and treated as in Fig.1a. Splenocytes were analyzed on termination. Cells were stimulated with ex vivo with either HIV gag peptides (e-i) or anti-CD3/CD28 antibodies (k-n) for cytokine response detection. (a) High-dimensional UMAP plots characterize CD8+ T cells similarity between different groups based on cellular markers (CD45RA, CCR7, CD28, CXCR5, PD-1, CD57, KLRB1). (b) The distribution of CXCR5+ TCM (CD45RA-CCR7+) on CD8 T UMAP space. (c) The percentage of CXCR5+ T_CM_ in CD8 T cells. (d) Description of the experiment. (e) Representative plots for IFN-γ, IL-2 and CD107a expression in CD8 T cells after peptides stimulation. (f) Summary data for IFN-γ expression in CD8 T cells. (g) Summary data for IL-2 expression in CD8 T cells. (h) Summary data for CD107a expression in CD8 T cells. (i) Summary data for the percentage of IFN-γ+IL-2+CD107a+ cell in CD8 T cells. (k) Summary data for IFN-γ expression in CD8 T cells. (l) Summary data for IL-2 expression in CD8 T cells. (m) Summary data for CD107a expression in CD8 T cells. (n) Summary data for the percentage of IFN-γ+IL-2+CD107a+ cell in CD8 T cells. Data shown is from two experiments. Bar represents mean value. * = p<0.05.

To investigate whether CD24-Fc treatment could improve anti-HIV T cell responses, we conducted ex vivo splenocyte stimulation with HIV-1 gag peptides and anti-CD3/CD28 antibodies (Fig. 3d). Cytokine analysis showed an enhanced viral-specific T cell response in CD24-Fc-treated splenocytes, with increased numbers of IFN-γ or IL-2-producing cells compared to the cART + Ig group (Fig. 3e-g). Additionally, CD8+ T cells from CD24-Fc-treated animals exhibited higher CD107a expression, indicating greater cytotoxic potential in HIV-1-specific CD8+ T cells (Fig. 3e and 3h). HIV-specific CD8+ T cells in PWH often exhibit reduced polyfunctionality, correlated with the failure of viral control [52, 53]. To identify HIV-1-specific polyfunctional T cells, we assessed IFN-γ, IL-2, and CD107a triple-positive cells upon viral stimulation. The frequency of these polyfunctional HIV-1-reactive CD8+ T cells was significantly higher in the CD24-Fc group than in the cART + Ig group (Fig. 3i). General T cell functionality also improved with CD24-Fc, as shown by increased cytokine production and polyfunctional T cells following anti-CD3/CD28 stimulation (Fig. 3k-n). These results suggest that CD24-Fc treatment substantially improves T cell functionality and anti-HIV responses in vivo, correlated with the restoration of CXCR5+ CD8 T_CM_ cells during suppressive cART.

### CD24-Fc treatment delays HIV-1 rebound and reduces viral pathogenesis after cART cessation

We hypothesized that the improved anti-HIV T cell response observed with CD24-Fc treatment during suppressive cART might promote a reduction in the HIV reservoir and delay viral rebound after cART withdrawal. To test this, we assessed cell-associated HIV-1 RNA and DNA levels in splenocytes from prior experiments. Surprisingly, there was no observed difference in cell-associated HIV-1 RNA or DNA between the cART + Ig and cART + CD24-Fc groups (Fig. 4a and 4b). However, CD24-Fc treatment reduced the cell-associated HIV-1 RNA/DNA ratio compared to the cART + Ig group, indicating that viral gene expression in reservoir cells was very low during CD24-Fc treatment with cART (Fig. 4c). Interestingly, the frequency of CXCR5+ CD8 T_CM_ cells was negatively correlated with the HIV-1 RNA/DNA ratio in splenocytes, suggesting that CXCR5+ T_CM_ cells may contribute to restricting viral transcription (Fig. 4d). To explore if CD24-Fc treatment could delay HIV-1 rebound, we conducted an additional experiment as shown in Fig. 1, stopping cART eight days after the final CD24-Fc injection to exclude any direct effect of CD24-Fc on viral rebound. Viral load analysis showed a one-week delay in viral rebound in the CD24-Fc group compared to the Ig group (Fig. 4e). All animals were euthanized at week 18 when viremia stabilized in all subjects. No significant differences in cell number (Fig. S2a-c) or T cell activation between CD24-Fc and Ig-treated groups were observed upon analysis of splenocytes (Fig. S2c).

**Figure 4.**
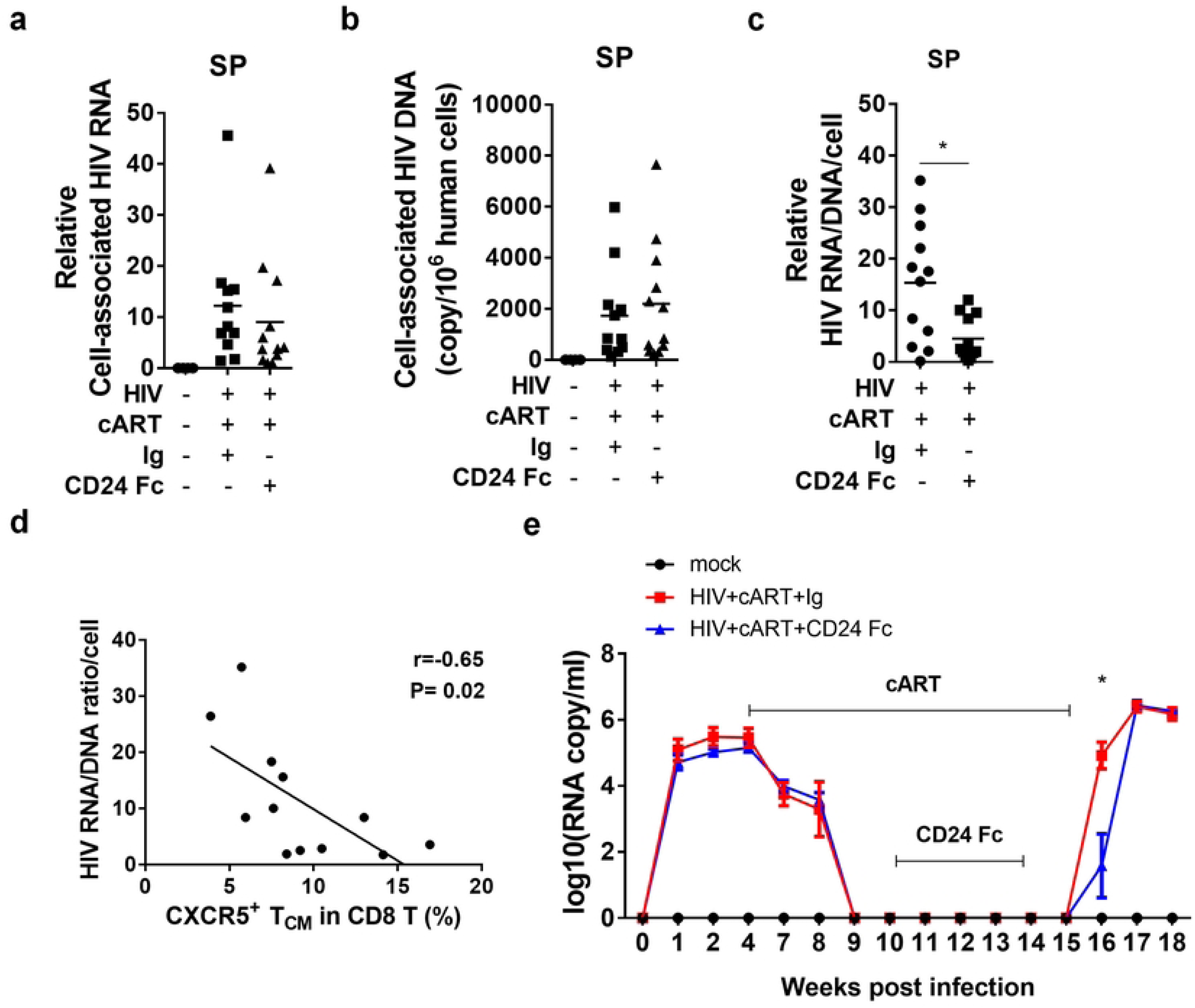
CD24-Fc treatment delays HIV-1 rebound after cART cessation. Humanized mice were infected with HIV treated as in Fig.1a. (a) The level of cell associated HIV RNA in splenocytes. (b) The copy number of cell-associated HIV-1 DNA per million hCD45+ cells. (c) The ratio of HIV cell-associated RNA and DNA at per cell basis. (d) Correlation between the ratio of HIV cell associated RNA/DNA and the frequency of CXCR5+ PD-1+ central memory CD8 T cell. (d) Plasma HIV viral load in a separate experiment with HIV rebound after cART cessation. Bar represents mean value. * = p<0.05. Error bar indicates mean value ±s.e.m.

Since viral reservoirs were not reactivated during CD24-Fc and cART treatment, reservoir cells likely evaded targeting by HIV-specific cytotoxic lymphocytes (CTLs). To enhance therapeutic efficacy, we added two doses of poly (I:C), known to both reactivate HIV-1 replication and enhance anti-HIV immunity, during CD24-Fc and cART treatments. Additionally, we extended CD24-Fc treatment post-cART cessation until the experiment’s end to leverage its anti-inflammatory effects after HIV rebound (Fig. 5a). Remarkably, in the CD24-Fc + poly (I:C) group, HIV viremia rebound was delayed by approximately two weeks compared to the cART + Ig group. Viremia levels in all infected mice reached a similar point three weeks post-cART withdrawal, at which point all animals were euthanized (Fig. 5a). Upon termination, the CD24-Fc + poly (I:C) group showed a higher CD4+/CD8+ T cell ratio compared to the cART + Ig group, which only partially restored the CD4/CD8 ratio relative to mock and HIV-only groups (Fig. 5b and 5c). Additionally, CD24-Fc + poly (I:C) treatment decreased the frequency of activated (HLA-DR+CD38+) CD8 T cells compared to the cART + Ig group (Fig. 5d and 5e). No significant differences in human leukocyte and T cell numbers in the spleen were noted between cART + Ig and CD24-Fc + poly (I:C) groups (Fig. S3). In summary, CD24-Fc treatment provided multiple benefits, including resolution of HIV-1 immune pathogenesis, enhanced anti-HIV T cell responses during suppressive cART, and delayed viral rebound following cART cessation in humanized mice.

**Figure 5.**
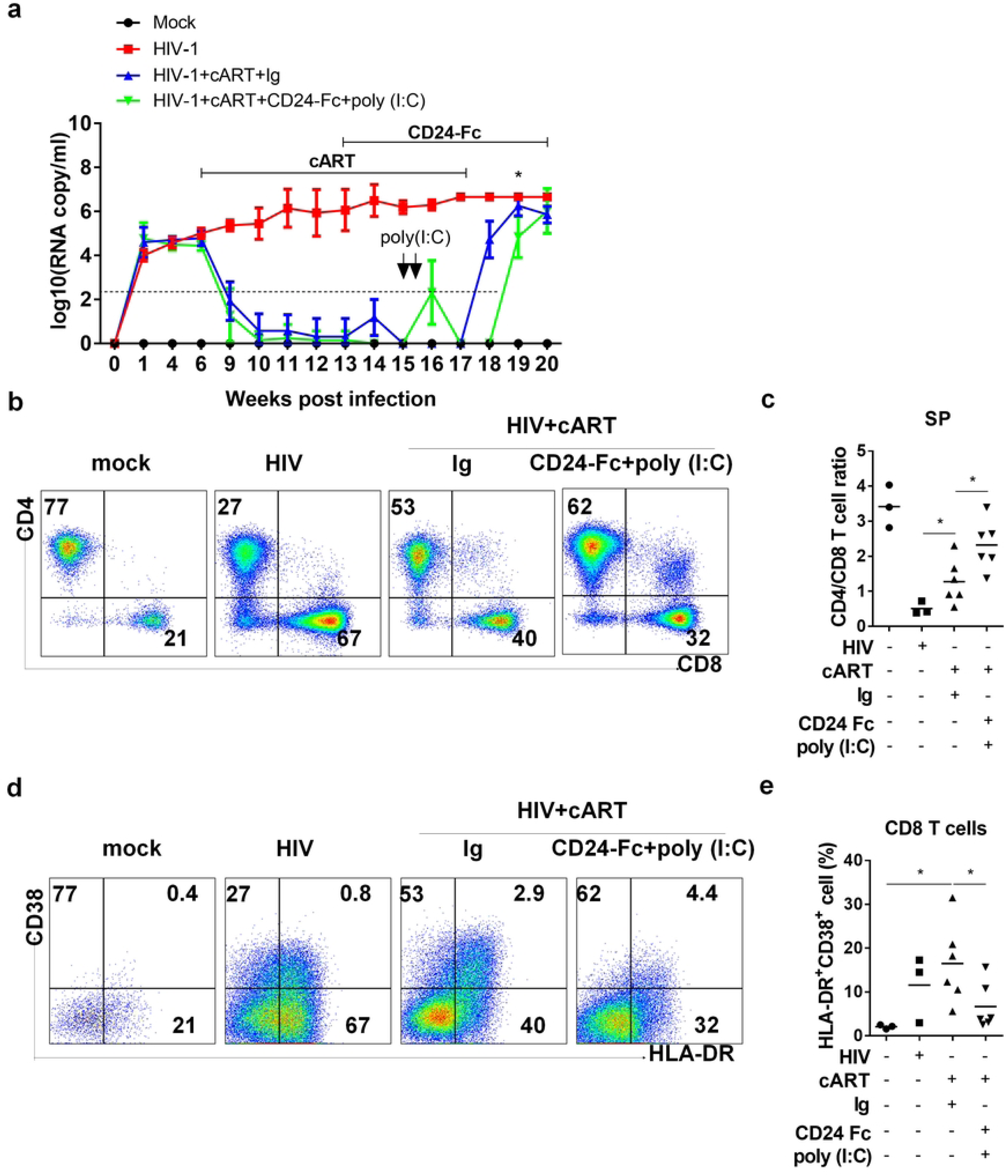
Sustained CD24-Fc treatment plus poly (I:C) further delay HIV-1 rebound and reduce HIV pathogenesis after cART cessation. Humanized mice were infected with HIV and then administered with cART in daily diet starting at 6 wpi and stopped at 17wpi. CD24-Fc treatment introduced at 13 wpi through i.p injection twice a week until termination. Ploy(I:C) was treated by one i.p. injection in two continuous days starting at 15 wpi. All animals were terminated at 20 wpi. Cells from spleen were analyzed by flow cytometry. (a) Viremia detected at the indicated time. (b) Representative FACS plots show the frequency of CD4+ and CD8+ T cell in CD3+ cells. (c) Summary graph for CD4 and CD8 T cell ratio. (d) Representative FACS plots show the frequency of HLA-DR+ and CD38+ cells in CD8+ T cells. (e) The percentage of HLA-DR and CD38 double positive CD8 T cells. Bar represents mean value. * = p<0.05. Error bar indicates mean value ±s.e.m.

### CD24-Fc treatment increases CXCR5 expression and functionality in CD8 T cells in PWH PBMCs in vitro

To evaluate the effects of CD24-Fc on immune cells from PWH, we cultured peripheral blood mononuclear cells (PBMCs) from both HIV-negative donors (healthy controls, HC) and virologically suppressed PWH. Cells were treated in vitro with or without CD24-Fc protein for 9 days, then stimulated with anti-CD3/CD28 antibodies to assess T cell functionality (Fig. 6a). Initially, we compared the number and function of T cells from HC and PWH after nine days of culture with IL-2 and IL-7. CD8 T cells from HC were more viable than those from PWH and produced more IFN-γ or TNF-α following stimulation with anti-CD3/CD28 antibodies (Fig. S4a and S4b). Examining polyfunctional CD8 T cells (those producing IFN-γ, TNF-α, granzyme B, and CD107a simultaneously; Fig. S4c and S4d), we observed that PWH cells showed fewer polyfunctional CD8 T cells than HC cells (Fig. S4e). However, CD24-Fc treatment appeared to mitigate this deficit, increasing the overall number of CD8 T cells in PWH cultures relative to controls (Fig. 6b). CD24-Fc treatment also increased the number of CXCR5+ memory CD8 T cells, though their frequency in CD8+ T cell population remained unchanged (Fig. 6c, Fig. 6d, Fig. S5a). In PWH cultures, CD24-Fc treatment led to an increase in the number and frequency of CD8 T cells expressing effector markers such as IFN-γ, TNF-α, granzyme B, and CD107a, compared to control groups (Fig. S5b and S5c). However, no difference in expression intensity was observed (Fig. S5d). Using UMAP clustering, we further analyzed polyfunctional CD8 T cells expressing all four effector markers and found that while their frequency remained constant (Fig. S5e), the number of polyfunctional CD8 T cells increased significantly with CD24-Fc treatment (Fig. 6g). Interestingly, upon stimulation through TCR signaling, CD8+ T cells from PWH PBMCs expressing cytokines are predominantly CXCR5+ memory T cells, particularly CXCR5+ T_CM_ (Fig. S6). Thus, the presence of CXCR5-expressing memory CD8+ T cells appears to correlate with T cell response in PWH. These findings suggest that CD24-Fc treatment improves CD8 T cell viability and enhances the function of CD8 T cells in PWH, indicating its potential role in protecting and supporting the functionality of these critical immune cells during chronic HIV infection.

**Figure 6.**
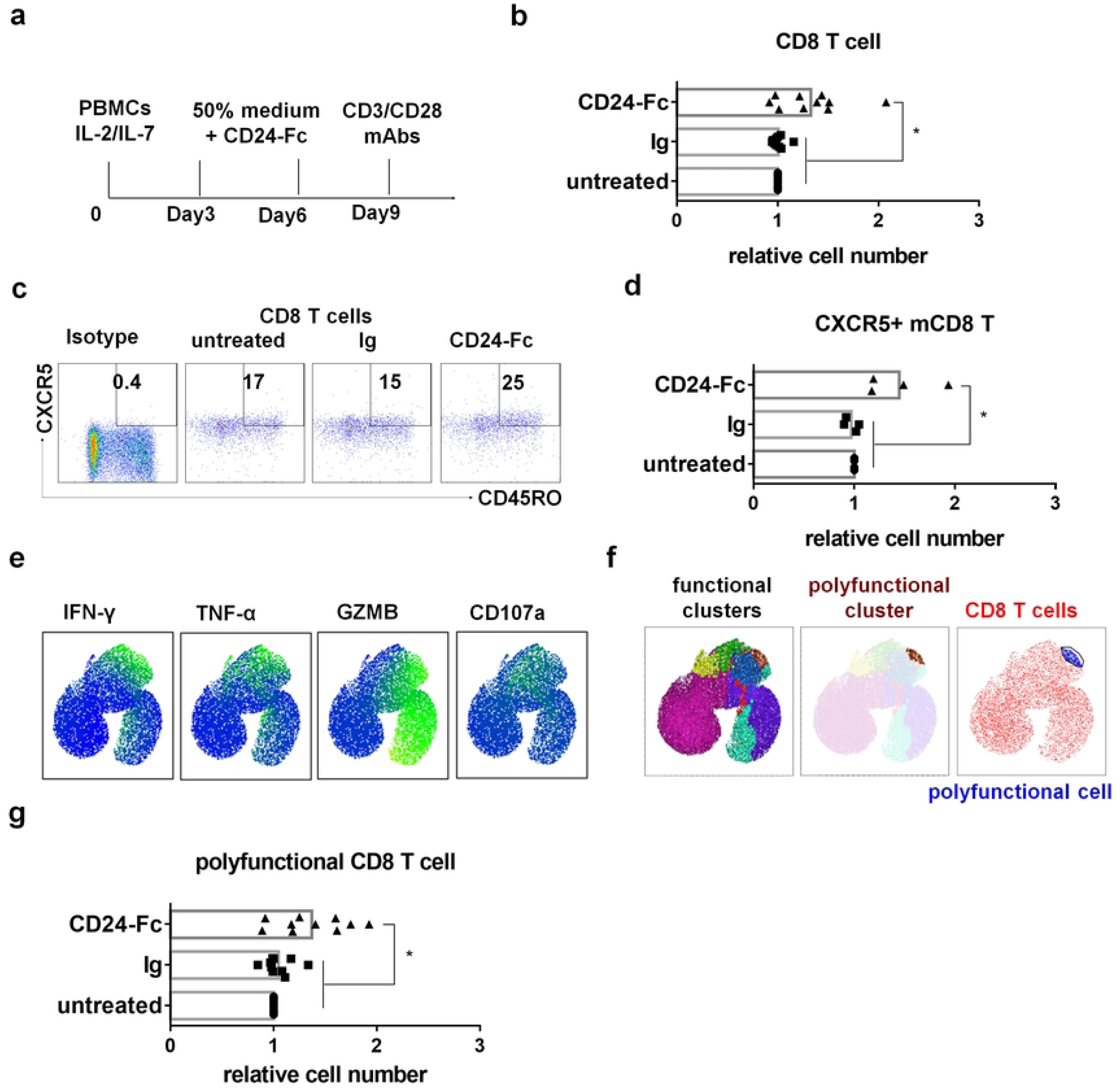
CD24-Fc treatment increases CXCR5+PD-1+ memory T cell and T cell functionality in PWH PBMCs in vitro. PBMCs from PWH were cultured with CD24-Fc for 9 days and stimulated with anti-CD3/CD28 antibodies on day 9 after culture. (a) Schematic description of experiment. (b) The relative number of CD8 T cell number in culture on day 9. (c) Representative FACS plots show the frequency of CXCR5+ memory cell in CD8 T cells. (d) Summary graph for the relative number of CXCR5+ memory CD8 T on day 9. (e) Different effector markers expression in CD8 T cells UMAP space. (f) Polyfunctional cluster identified by FlowSOM (brown) and polyfunctional cells gated on accordingly (blue). (g) The relative number of polyfunctional CD8 T cell in individual donor in different groups. Data shown is from two experiments. Bar represents mean value. * = p<0.05.

## Discussion

Preclinical research on CD24-Fc in SIV-infected primates suggests it can reduce inflammation and slow AIDS progression, showing promise for managing chronic immune activation in HIV and SIV infections[32, 33]. This study reveals that CD24-Fc therapy, when combined with cART, significantly reduces inflammation and immune activation in HIV-1 infected humanized mice. CD24-Fc appears to suppress HIV-1 LTR (long terminal repeat) activity, and notably, there were no viremia blips or viral reservoir reactivations observed in the treated mice. These findings suggest that CD24-Fc therapy might promote quiescence of viral replication within reservoir cells, thus helping them evade detection by immune surveillance. This ability to maintain reservoirs in a hidden state could make CD24-Fc an attractive adjunct to cART in controlling chronic HIV infection. The study proposes a follow-up experiment to confirm CD24’s role in this mechanism by blocking CD24 in vivo during cART to observe whether reservoir activation changes, potentially validating CD24-Fc’s ability to suppress reservoir activation during cART[54].

The specific cell types targeted by CD24-Fc in this study remain unclear. CD24 is known to primarily interact with Siglec-10 to deliver inhibitory signals that reduce inflammation[29]. In humans, Siglec-10 is predominantly expressed on dendritic cells, natural killer (NK) cells, and B cell[55]. Research also indicates that activated CD4+ T cells upregulate Siglec-10 expression[56], which is associated with persistent abnormal CD4 T cell activation PWH despite viral suppression by cART.[57]. In this study, CD24-Fc treatment appeared to enhance CD4 T cell recovery in vivo. This improvement may arise either from a direct effect on CD4 T cells or indirectly via systemic inflammation reduction. However, experimental data suggest that CD24-Fc does not directly influence HIV replication in purified activated CD4 T cells, nor does it impact CD4 T cell counts in vitro. Therefore, the ability of CD24-Fc to suppress HIV replication and improve CD4 T cell levels in vivo seems to function through an indirect pathway, likely mediated by its broader anti-inflammatory effects.

This study demonstrated that CD24-Fc treatment can restore CXCR5+ CD8 memory T cells, which are crucial for anti-viral immunity, correlating with lower HIV viremia levels before antiretroviral therapy[51]. Consistent with previous findings, we observed that restoration of CXCR5+ CD8 memory T cells are associated with suppressed viral replication and enhanced T cell polyfunctionality in humanized mice in vivo and in PBMC cultures from PWH in vitro. Based on these findings, we hypothesize that a combined immunotherapy involving CD24-Fc, PD-1/PD-L1 checkpoint inhibitors, and latency-reversing agents (LRAs) could further boost the anti-HIV immune response. This approach could potentially reduce the HIV reservoir and delay viral rebound upon cART cessation. This combination therapy will be investigated in future studies.

One possible disadvantage of CD24-Fc treatment is that macrophages phagocytosis can be blocked by CD24-Fc treatment when it binds to Siglec-10 on macrophages as the recent reports showed that CD24 exerts a novel ‘don’t eat me’ signal in tumor cells [54]. Thus, an evaluation of CD24-Fc therapeutic potential in non-human primate model with the functional intact myeloid lineage cells should be done in future.

A potential drawback of CD24-Fc treatment is its possible interference with macrophage phagocytosis. CD24-Fc binding to Siglec-10 on macrophages might activate a “don’t eat me” signal, a mechanism recently observed in tumor cells involving CD24, which could impede macrophage activity. This raises concerns about its effect on immune clearance functions.

Therefore, an evaluation of CD24-Fc therapeutic potential in non-human primate models with an intact and functional myeloid lineage is essential. Such studies would provide insight into the balance between its anti-inflammatory benefits and possible limitations related to immune cell phagocytosis, guiding the safe development of CD24-Fc as a treatment option.

## Material and methods

### Ethics statement

All animal experiments were reviewed and approved by the Institutional Animal Care and Use Committee (IACUC) at the University of North Carolina at Chapel Hill (Protocol ID: 16-073).

### Humanized mice

Humanized mice were generated as previously described [17, 42, 58–61]. Briefly, NOD-Rag1^null^IL2rg^null^ (NRG) neonates (1-to-5 days old) were irradiated (250 rads) and injected with 2 x 10⁵ human CD34+ hematopoietic stem cells (HSCs) into the liver. HSCs were isolated from human fetal liver tissues obtained from elective or medically indicated pregnancy terminations through a non-profit intermediary working with outpatient clinics (Advanced Bioscience Resources).

### HIV-1 infection of humanized mice

Humanized mice were infected via retro-orbital injection with HIV-1_JRCSF_ stocks (10 ng p24/mouse) or 293T mock transfection supernatants for control mice.

### cART regimens in humanized mice

Individual tablets of TRUVADA (tenofovir/emtricitabine; Gilead Sciences) or raltegravir (Merck) were crushed into fine powder and manufactured as 5BXL by TestDiet based on previously published [15, 37, 62].

### CD24-Fc Fusion Protein Treatment

CD24-Fc protein was obtained from Yang Liu’s lab as a gift. The recombinant CD24-Fc fusion protein and IgG-Fc was manufactured according to current ideal manufacturing procedures as previously described [32]. Humanized mice were administered twice a week with 200μg recombinant protein each dosage through intraperitoneal injection (i.p.).

### HIV viral load in plasma

Blood was collected by tail vein bleeding using EDTA as an anticoagulant, and plasma was stored at −80℃ until assay. HIV-1 RNA was extracted from plasma using the Viral RNA Mini Kit (Qiagen) and quantified by real-time PCR with the TaqMan® Fast Virus 1-Step PCR kit (ThermoFisher Scientific) on a QuantStudio 6 Flex PCR system (Applied Biosystems), with a detection limit of 400 copies/ml [15, 17, 37, 42].

### Real-time PCR

For detecting interferon-stimulated genes (ISGs), RNA was isolated from splenocytes using the RNeasy Plus extraction kit (Qiagen) and converted to cDNA using SuperScript III First-Strand Synthesis (Invitrogen). ISG levels in cDNA were quantified by real-time PCR with human gene-specific primers as previously described [15, 16].

For cell-associated HIV-1 DNA, nucleic acid was extracted from cells or tissues using DNeasy mini kit (Qiagen). HIV-1 DNA was quantified by real-time PCR. Genomic DNA of ACH2, which contains one copy of HIV genome in each cell, was serially diluted in mouse leukocytes DNA to generate a standard curve [15].

For cell-associated HIV-1 RNA, RNA was extracted from cells or tissues using RNeasy plus mini kit (Qiagen). HIV-1 RNA was detected as previously described[15, 17, 42]. The HIV-1 gag RNA expression was normalized to human CD4 mRNA level and relative HIV-1 gene expression levels were calculated according to 2^-ΔΔCT^ [15, 63].

### Anti-HIV T cells detection

Cells from humanized mice spleen were stimulated ex vivo with an HIV gag peptide pool (2 μg/ml per peptide; PepMix HIV (GAG) Ultra, JPT Innovation Peptide Solutions) and human CD28 antibody (2 μg/ml) for 3 hours without, and then 5 hours with, brefeldin A. Cells were fixed, permeabilized, and subjected to intracellular staining.

### Participants

Ten HIV-infected participants on suppressive cART were recruited from the labs of Nilu Goonetilleke (UNC Chapel Hill), R. Brad Jones (Cornell University), and Poonam Mathur (University of Maryland, Baltimore) (Table S1). Participants had plasma HIV RNA levels ≤ 40 copies/ml, as measured by the Abbott Real-Time HIV-1 PCR at the time of sample collection, and all had been on cART for at least 12 months. Blood was collected by standard venipuncture, and leukapheresis was performed to obtain peripheral blood mononuclear cells (PBMCs). Written informed consent was obtained from all participants under an approved IRB protocol.

### In vitro human PBMC assays

PBMCs from PWH or HIV-negative donors were cultured at 1 x 10⁶ cells/ml in 10% FBS RPMI-1640 containing 20 U/ml IL-2, 10 ng/ml IL-7, and CD24-Fc protein (10 μg/ml) or IgG (10 μg/ml). Controls received no treatment. Every 3 days, 50% of the medium was replaced with fresh medium containing 40 U/ml IL-2, 20 ng/ml IL-7, and 20 μg/ml CD24-Fc protein. On day 9, live cells were counted and cultured in complete medium with anti-CD3 (1 μg/ml, clone 30-F11; Biolegend) and anti-CD28 (1 μg/ml, clone CD28.2; Biolegend) antibodies. Cells were stained for flow cytometry after 6 hours of incubation with brefeldin A at 37°C.

### Flow cytometry and data analysis

For intracellular staining, cells were stained with surface markers first, and then permeabilized with cytofix/cytoperm buffer (BD Bioscience, cat#554714), followed by intracellular staining. Anti-human antibodies were purchased from Biolegend, including anti-CXCR5 (clone:J252D4), anti-CD4 (clone:RP4-T4), anti-CD8 (clone:HIT8a), anti-CD3 (clone:HIT3a), anti-CD45 (clone:H130), anti-CD45RA (clone:H100), anti-CCR7 (clone:G043H7), anti-KLRG1 (clone:SA231A2), anti-CD161 (clone:W18070C), anti-CD57 (clone:QA17A04), anti-HLA-DR (clone:L243), anti-CD38 (clone:HIT2), anti-PD-1 (clone:NAT105), anti-IFN-γ (clone:4S.B3), anti-TNF-α (clone: Mab11), granzyme B (clone: GB11), CD107a (H4A3) and anti-IL-2 (clone:MQ1-17H12). Anti–mouse CD45 (clone: HI30) and LIVE/DEAD Fixable Aqua Dead Cell Stain Kit (cat#L34957) were purchased from Invitrogen. Flow cytometry was performed using BD LSRFortessa (BD Biosciences) and analyzed by FlowJo 10 (FLOWJO, LLC).

### UMAP and clustering analysis

FCS3.0 data files were imported into FlowJo software version 10.8.1 (FlowJo LLC). All samples were compensated electronically. Dimensionality reduction was performed using the UMAP. For UMAP analysis, live CD3+CD4-CD8+ populations were concatenated. UMAP plots were generated with default settings and excluding all parameters used upstream in the gating strategy (CD3, CD4 and CD8). The same numbers of CD8+ T cells from HIV negative and HIV positive individuals were similarly applied to UMAP analysis. Markers considered in data from humanized mice include CD45RA, CCR7, CXCR5, PD-1, CD28, CD57, CD161 and KLRG1. Markers considered in data from humanized mice include CD45RA, CCR7, PD-1, CD28, CD57, CD161 and KLRG1. Markers considered in data from humanized mice include CD45RA, CCR7, PD-1, CXCR5, PD-1, IFN-γ, IL-2, CD107a, granzyme B. We identify polyfunctional cluster expressing multi T cell effector markers using FlowSOM version 4.0.0 plugin. ClusterExplorer version 1.7.6 plugin was used to map clusters on UMAP plot and generate heatmap for individual cluster. To characterize cell cluster, manual gating was applied on UMAP space based on the auto-clusters, and then pseudo-colored was applied on the UMAP plot.

### Statistical analysis

Statistical analyses were conducted using unpaired 2-tailed Student’s t-tests and one-way ANOVA with Bonferroni multiple comparisons in GraphPad Prism (GraphPad Software, San Diego, CA). A p-value < 0.05 was considered statistically significant. Bars represent mean values, with error bars indicating ±standard error of the mean (s.e.m.).

## Acknowledgements

We thank Liquan Chi, Yaxu Wu, Weirong Yuan and Yichen Lu for their technical supports as well as all current and previous Su lab members for critical reading and/or discussion of the manuscript. We thank the University of North Carolina (UNC) Division of Laboratory Medicine for animal care; University of North Carolina at Chapel Hill Center for AIDS Research (P30 AI50410). We thank Flow Cytometry Core Facility at University of North Carolina at Chapel Hill. We thank the Flow Cytometry Shared Service of the University of Maryland Marlene and Stewart Greenebaum Comprehensive Cancer Center for core support. This study was supported in part by grants from the National Institutes of Health (AI080432, AI077454 and AI095097 to LS). The funders had no role in data analysis, preparation, or decision to publish the manuscript.

## Authorship

G.L. designed, performed experiments, prepared figures, analyzed data and wrote the paper; G.L., J.M., H.Y., R.T., Y.L., X.H. and M.F. performed the experiments; P.M. and S.K. provided PWH samples; P.Z and Y.L provided CD24-Fc protein; L.S. conceived the research project, planned, designed experiments, and wrote the paper. Current address correspondence to: H.Y., Institute of Infectious Diseases, Guangzhou Eighth People’s Hospital, Guangzhou Medical University, Guangdong, China. X.H., Departments of Geriatrics, The First Hospital of Jilin University, Jilin, China. M.F., Department of Gastroenterology, Kanazawa University Hospital, Kanazawa, Ishikawa, Japan.

